# Identification of Pain-Associated Effusion-Synovitis from Knee Magnetic Resonance Imaging by Deep Generative Networks

**DOI:** 10.1101/2023.05.04.539501

**Authors:** Pin-Hsun. Lian, Tzu-Yi Chuang, Yi-Hsuan Yen, Gary Han Chang

## Abstract

**Objectives:** To identify the source and location of osteoarthritis-induced pain symptoms, we used deep learning techniques to identify imaging abnormalities associated with pain from magnetic resonance imaging (MRI) of knees with symptoms of symptoms of osteoarthritis pain.

**Methods:** Pain-associated areas were detected from the difference between the MRI images of symptomatic knees and their respective counterfactual asymptomatic images generated by a Generative adversarial network. A total of 2,225 pairs of 3D MRI images were extracted from patients with unilateral pain symptoms in the baseline and follow-up cohorts of the Osteoarthritis Initiative. Subsequently, pain-associated effusion-synovitis were characterized into subregions (patellar, central, and posterior) using an anatomical segmentation model.

**Results:** We found that the volumes of pain-associated effusion-synovitis were more sensitive and reliable indicators of pain symptoms than the overall volumes in the central and posterior subregions (odds ratio [OR]:3.23 versus 1.77 in the central region, and 3.18 versus 2.66 in the posterior region for severe effusion-synovitis). For mild effusion-synovitis, only pain-associated volume was found to be associated with pain symptoms, but not with overall volume. Patients with significant pain-associated effusion-synovitis in the patellar subregion had the highest increased odds of pain symptoms (OR=4.86).

**Conclusion:** To the best of our knowledge, this is the first study to utilize deep-learning-based models for the detection and characterization of pain-associated imaging abnormalities. The developed algorithm can help identifying the source and location of pain symptoms and in designing targeted and individualized treatment regimens.

## 1. Introduction

Osteoarthritis (OA) is the most prevalent, debilitating, and age-related disease worldwide (1). Advanced knee OA is often accompanied by progressive worsening of pain, reduced mobility, and disability (2). As one of the main causes of disability globally, the burden of knee OA is expected to increase due to aging of the population. Knee pain with osteoarthritis can be accompanied by a wide variety of imaging findings, such as effusion-synovitis, bone marrow lesions (BMLs), and cartilage defects (3). However, there is a well-known disparity between the imaging findings and OA pain symptoms (4,5). Investigating the association between pain symptoms and imaging characteristics of the knees is further complicated by subjective and non-structural confounding factors that cannot be observed on knee imaging (5–7). Neuropathic mechanisms, depression, history of injuries, and genetic predisposition can all contribute to the symptoms of OA pain. As a result, it remains challenging to pinpoint the source and location of imaging findings associated with pain symptoms, hindering the development of targeted and localized treatment strategies, such as unicondylar replacements and cryoneuroablation procedures (8). To minimize the influence of personal confounders, an effective approach is to focus on patients with unilateral knee pain symptoms (9). By comparing the knee with pain symptoms with the contralateral normal knee in a patient with unilateral symptoms, the influences of structural findings were easier to isolate and eliminate from individual confounders. This approach has previously been used to demonstrate the association between radiographic OA and knee pain to identify regions of roughness that were more likely to contribute to knee pain on magnetic resonance imaging (MRI) (10) and to study focal knee lesions in patients with unilateral symptoms (10).

Magnetic resonance imaging (MRI) has been able to reveal more consistent connections between imaging findings and OA-induced pain symptoms in different studies (11,12). However, as MRI findings suspected to be associated with pain symptoms often appear in multiple knee locations, and asymptomatic knees can also present with these MRI findings, studying the association between imaging abnormalities and pain symptoms via MRI scoring is a time-consuming and expertise-demanding process. Comparing MRI findings between knees traditionally depends on various types of semiquantitative scoring systems implemented by expert readers (13–15), and studies often only select a subset of data for expert reading. Additionally, MRI scoring criteria are susceptible to uncertainty due to the potential for disagreement among experts regarding severity thresholds (16). Although several types of inflammatory lesions have been proposed to be associated with knee pain, such as synovitis and bone marrow lesions (BMLs) (17–19), identifying and quantifying the source and location of the MRI findings associated with pain symptoms remains difficult. Therefore, the development of an automatic approach to facilitate the identification of pain-associated MRI abnormalities is of great interest. Accurate identification and qualification of imaging findings associated with OA symptoms may help identify candidates for targeted therapies.

In recent years, the rapid advancement in deep learning techniques has enabled their use in a range of medical imaging applications (20,21), including the automatic diagnosis, segmentation, and progression prediction of knee radiography and MRIs (22–28). In particular, deep generative models have shown the ability to create high-fidelity and realistic images and perform anomaly detection in a wide variety of biomedical imaging modalities including computed tomography, MRI, radiography, and microscopy (29,30). In this study, we investigated the ability of deep learning techniques to automatically characterize and quantify imaging abnormalities associated with pain by studying patients with unilateral knee pain. We utilized generative adversarial networks (GANs) to quantify the extent of two types of MRI findings, BMLs and effusion-synovitis, as their correlation with OA pain symptoms has been shown in large cohort studies (12). This was achieved by utilizing a GAN that created asymptomatic counterfactual images of a knee from an originally symptomatic knee with which it shared the same anatomical structures. An asymptomatic contralateral knee of the same patient was used as the reference point. We studied sagittal intermediate-weighted turbo spin-echo (IW-TSE) MRI images in which both types of inflammatory lesions (effusion-synovitis and BMLs) were highly visible. Regions with abnormalities associated with pain were identified by calculating the difference between the original MR image and the generated asymptomatic counterfactual image.

We further quantified the extent of BMLs and effusion-synovitis in individual subregions with an anatomical segmentation network, which we trained by transferring structural information from sagittal double-echo steady state (DESS) MR images to IW-TSE images using a domain adaptation technique. We then quantified the distribution of pain-associated BMLs and effusion-synovitis in individual subregions to differentiate how imaging abnormalities in each subregion may contribute differently to pain symptoms. Finally, we demonstrated that the abnormal areas identified by our approach captured the essence of the symptoms-related features. Specifically, we found that an independent model trained exclusively on the identified areas achieved comparable predictive performance to models trained on full MR images in predicting pain symptoms. The main contributions of our work are:

1. We demonstrate, for the first time, the ability to automatically detect pain-associated abnormal areas in MRI images of OA knees using adversarial generative networks.
2. We introduce a novel approach for incorporating Monte-Carlo dropout and uncertainty estimation into anomaly detection, which aids in the identification of abnormal areas that are highly likely to contribute to pain symptoms.
3. By characterizing the detected abnormal regions into subregions by a cross-modality segmentation model, we showed that effusion-synovitis in the proximity of patella exhibits the highest likelihood of contributing to OA symptoms. This finding is consistent with previous studies in the literature and underscores the clinical relevance of the present study.

## 2. Methods

### 2.1 Model structure overview

The proposed framework for the automatic identification of pain-associated abnormalities is summarized in **Figure 1A**. The generative network (G_pain_, **Figure 1A**) created an ensemble of counterfactual images without pain symptoms from the original MR images of knees with pain symptoms by using the contralateral asymptomatic knee as the reference. Pain-associated regions were defined as areas where the generated counterfactual images exhibited a statistically significant difference from the original images, which were derived from the mean and variation of the pixel-wise difference between the counterfactual and original images **(Figure 1B)**. Three-dimensional volumes of sagittal IW-TSE MR images of paired knees from patients with unilateral knee pain were extracted from the Osteoarthritis Initiative (OAI) cohorts and cross-validated by five folds with mutually exclusive individuals. The training set was used to train the counterfactual generative model, which performed image-to-image translation from the symptomatic knee to the contralateral asymptomatic knee from the same patient. Subsequently, the identified abnormal regions were characterized as subregions of effusion-synovitis using an anatomical segmentation model (G_seg_, **Figure 1A**) that divided the MR images into subregions of the bone of the femur and tibia and subregions of the patellar, central and posterior effusion-synovitis. Finally, to investigate whether the associations between knee pain symptoms and MR imaging were sufficiently represented by the identified abnormal regions, we trained an independent pain classification model exclusively by using the shapes and locations of the abnormal regions learned previously from the generative model.

**Figure 1.**
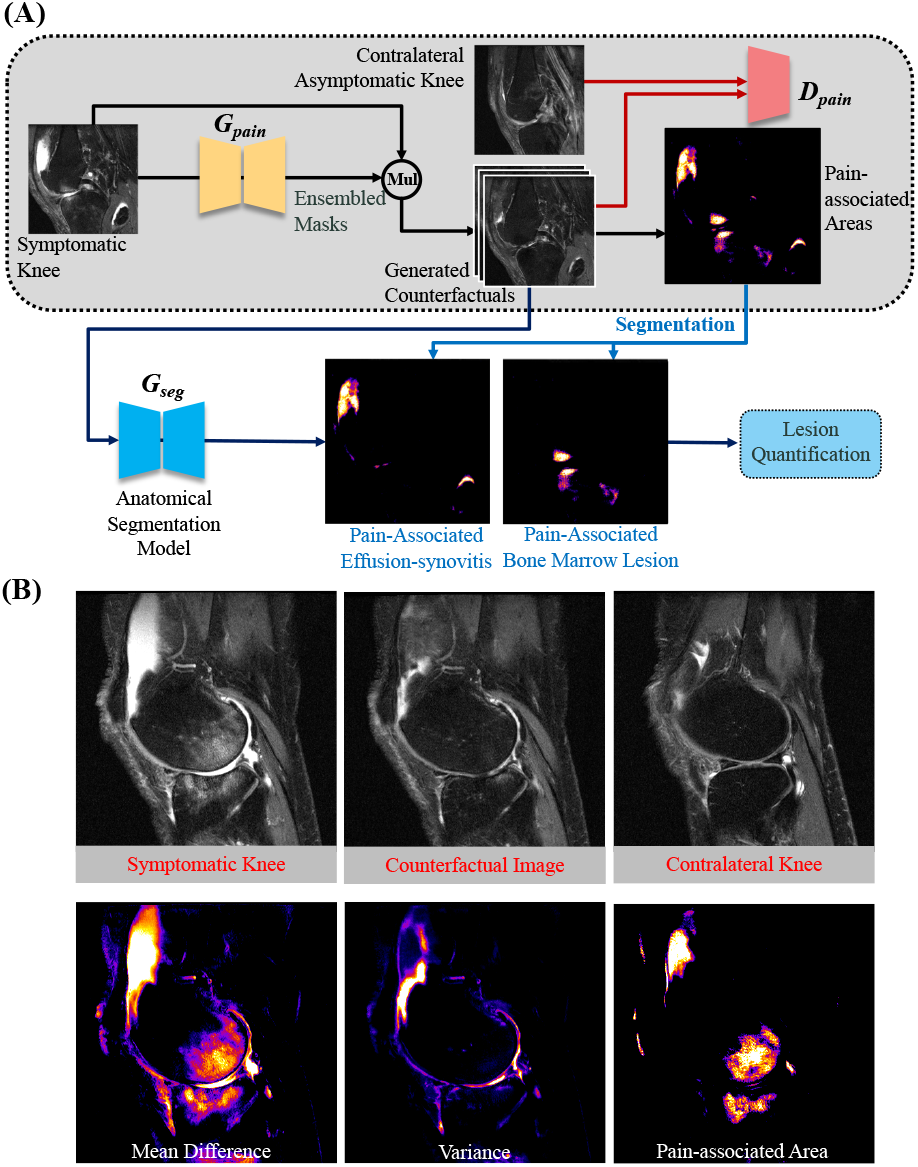
**Overview of the model framework. (A) The image of symptomatic knee is translated by the generative model, G_pain_, to synthesize the image of asymptomatic knee using the contralateral asymptomatic knee as the reference. The probabilistic generative model generated an ensemble of counterfactual images. The pain-associated abnormal areas were defined as the regions with statistically significant difference between original image to the ensembled counterfactual images. The detected regions were further segmented into the subregions of bone and effusion-synovitis by the segmentation model, G_seg_. (B) The pain-associated areas were defined first by calculating the mean and variation in differences between the original image to the ensemble of counterfactual images, then by identifying the areas from the counterfactual images that were statistically different than the original image.**

### 2.2 Imaging dataset and preprocessing

The imaging dataset for this study was acquired from the OAI baseline and follow-up cohorts (n = 4796). We selected patients who had significant contralateral differences in knee pain symptoms, defined as a Western Ontario and McMaster Universities Osteoarthritis Index pain score ≥ 4 between two knees, from among patients who had undergone sagittal IW-TSE imaging. In total, 2225 knee pairs met the criteria and were used for model construction and validation after excluding patients with imaging artifacts and metal implants. The selected knee pairs were separated into two splits, without overlapping individuals, for model training and validation. To facilitate a region-by-region comparison between each anatomical region from the knee pairs of the same patients, the images of the knee pairs were first aligned by the central MRI slice, defined as the slice with the most complete view of the posterior cruciate ligament. The images from the left knee were subsequently registered to the right knee images using affine transformation. An image stack of 11 slices adjacent to both the lateral and medial sides of the central slice was selected as the region of interest. The resulting image volumes had dimensions of 384 × 384 × 23 and their grayscale values were normalized to a range of 0 to 1. During the model training, the image volumes were randomly cropped in-plane to a size of 256 × 256 × 23.

### 2.3 Counterfactual generative network

The first step of this study was to detect abnormal regions associated with pain with high certainty. To this end, we used a counterfactual generation network (G_pain_ in **Figure 1A**) to only alter regions that were highly likely to be associated with pain, while leaving the details in other asymptomatic regions unchanged. Asymptomatic regions may include both healthy and low-grade lesions that are unlikely to contribute significantly to pain symptoms. To obtain the most likely counterfactual MR images without pain symptoms, as well as the associated uncertainty distribution, we employed a Bayesian GAN with Monte Carlo dropout, which generated an ensemble of plausible counterfactual MRI simultaneously for each input MRI. This approach has frequently been used to assess the uncertainties associated with medical imaging segmentation (31,32). By eliminating regions with high uncertainty in the counterfactuals generated, we can differentiate between pain-associated abnormal regions and those that may contain low-grade asymptomatic lesions.

To help preserve the pixel-level details of the input MR image, a counterfactual generation network was designed based on the frequently utilized U-shaped structures (33), which contain direct connections between the encoding and decoding layers of the network. These connections help transfer pixel-level information, such as details of healthy tissues, to the generated image. Similar architectures have been frequently utilized in high-definition image translation tasks in medical imaging. Furthermore, we utilized a residual learning approach, wherein the model created a counterfactual image through a combination of an original image and a generated mask whose values ranged from 0 to 1. This encourages the model to keep irrelevant regions intact while focusing exclusively on regions associated with pain symptoms.

During the training of the counterfactual generative model, an MRI volume of the knee showing pain symptoms was taken as the input, and the generated counterfactual knee was compared to the contralateral knee without pain symptoms simultaneously by the discriminator (D_pain_ in **Figure 1A**), which attempted to play against the generator by differentiating the real and counterfactual asymptomatic images. This adversarial training scheme ensured that the generated images maintained an imaging quality indistinguishable from the referenced images to synthesize high-fidelity MR images. The discriminator was also assisted by an auxiliary module whose task was to distinguish real MRI volumes of the knees with or without pain symptoms.

To detect the abnormal regions and associated uncertainties, a distribution of counterfactual images was generated by inferring the generative model multiple times (n=100 was chosen to ensure convergence of the results), while the average and uncertainties of the reduced intensities of each pixel were calculated over all the inferred samples. Finally, we identified abnormal regions where the intensity values were confidently reduced by the model, whereas regions with high uncertainties were excluded from the identified regions (**Figure 1B**).

### 2.4 Characterizing pain-associated abnormal areas by source and location

To characterize and quantify abnormal regions identified in our study, we used a deep neural network-based segmentation model to differentiate between effusion synovitis and bone marrow lesions. Specifically, we applied this model to IW-TSE MR images to automatically identify and segment (**G**_**effusion**_, **Figure 2A**) effusion-synovitis subregions in the patellar, central, and posterior regions of the knee joint, while excluding non-effusion regions such as bursitis. This approach, similar to that described in a previous study using non-contrast enhanced MRI to study effusion-synovitis (34), allowed us to characterize the effusion-synovitis subregions present in the studied sample more accurately.

**Figure 2.**
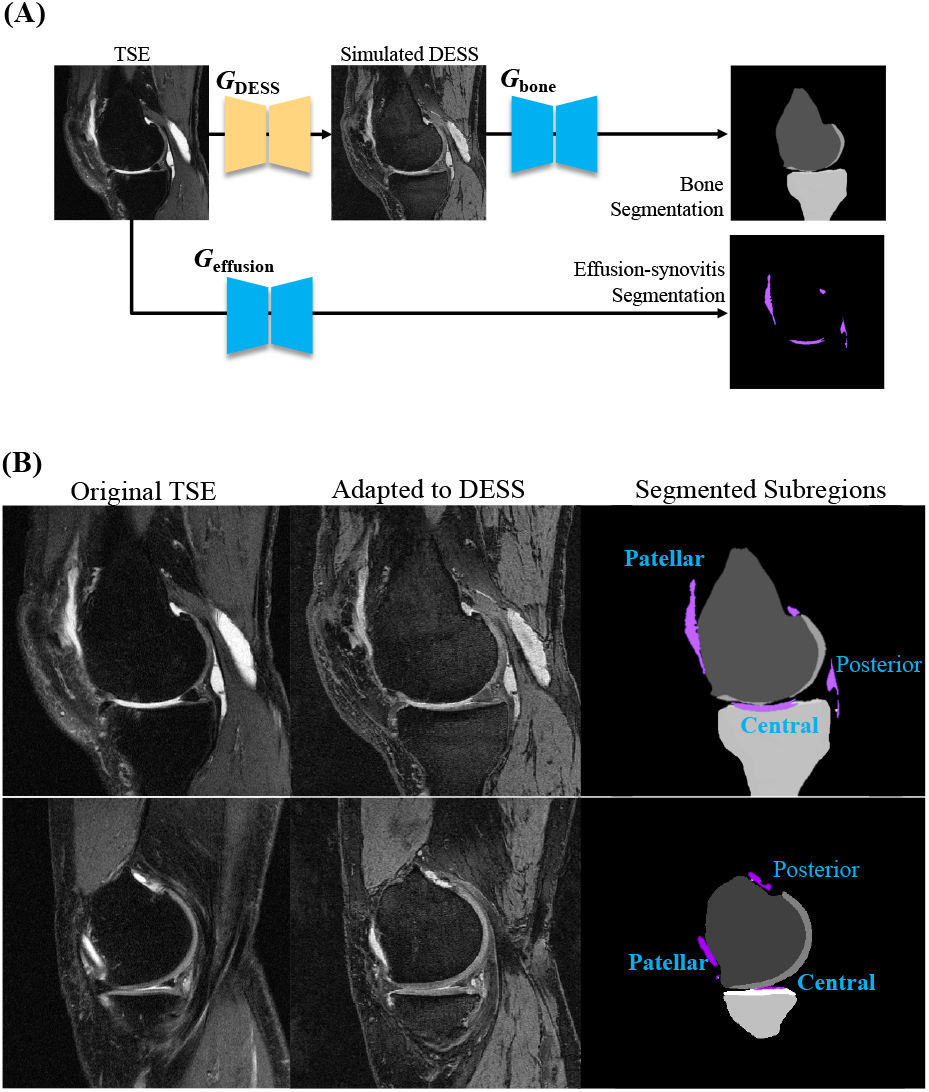
**Domain adaptation and segmentation models for characterizing the pain-associated areas into subregions. The TSE images were first adapted by a domain adaptation model which translated the TSE images to double-echo steady state (DESS) images. The adapted images demonstrated enhanced contrast between bone and cartilages, and were further segmented by a DESS segmentation model into subregions of patellar, central, and posterior effusion-synovitis, and bone subregions of femur and tibia. The segmented subregions were subsequently used to characterize the identified pain-associated areas.**

To characterize the area of the bone, as the IW-TSE MR images did not provide clear definition between bones, cartilages, and other anatomical structures, we applied domain adaptation method (35) to translate the IW-TSE images to DESS images. The domain-adaptation model (**GDESS, Figure 2A**) was trained with 6095 paired 2D slices of TSE and DESS images from the same knees that were manually selected and registered from the OAI. The counterfactual IW-TSE images generated previously were subsequently translated to visually resemble the DESS images and showed improved definitions of the bone, cartilage, and other structures (**Figure 2B**). The transformed images subsequently had their bone and cartilage structures automatically segmented using the DESS segmentation model (**G**_**bone**_, **Figure 2A**), which we previously developed to study bone and cartilage morphologies because of their superior definition of bone and cartilage in DESS images (36).

Since many BMLs are in close proximity to the subchondral bone plates, subtle differences between the edge of the bone and the surrounding cartilage, effusion, and other structures need to be differentiated accurately to allow proper identification of the BMLs. Therefore, we adjusted our bone segmentation model by taking advantage of the aforementioned Monte Carlo dropout model, which simultaneously generates bone segmentations and associated uncertainties from the generated counterfactual image. We subsequently excluded areas of high uncertainty in the segmented bone images to prevent hyperintensities, such as effusions, cartilage, and other non-bone structures from being misclassified as BMLs and negatively influence the characterization of pain-associated bone marrow lesions.

Finally, we studied the clinical relevance of the identified pain-associated areas by calculating the odds ratios (ORs) by comparing the odds of developing pain symptoms in each quartile using the first quartile as a reference for comparison. The ORs, their significance, and 95% confidence intervals (CI) were calculated using Fisher’s exact test. ORs were calculated using a 2 × 2 contingency table with binomial outcomes for pain symptoms.

## 3. RESULTS

### 3.1 Demography

Among the 2225 knee pairs from the OAI baseline and follow-up cohorts, the mean age was 61.1 ± 9.1 years at baseline and the mean BMI was 28.9 ± 4.7 kg/m^2^. Approximately 56.9% of the selected participants were women.

### 3.2 Generated counterfactual images

**Figure 3A** shows the MR images of knees with pain symptoms, their corresponding asymptomatic counterfactuals generated by the deep generative network, and the detected pain-associated areas. It can be observed that the generative model was able to differentiate anatomical subregions in details and altered exclusively on the subregions associated with pain. The model identified effusion-synovitis in the patellar region (red arrows) and infrapatellar fat pad (yellow arrows) as pain-associated. On the contrary, effusion-synovitis in the central area, not deemed pain-associated (green arrows), remains present in the images. In **Figure 3B**, the model showed the capacity to characterize distinct source of pain without prior knowledge of anatomical structures. It identified effusion-synovitis in the patellar regions as pain-associated (red arrows). Contrarily, the effusion-synovitis in the central region (green arrows) was not regarded as pain-associated, even when exhibiting lower intensity values and situated adjacent to the bone marrow lesions, which were considered pain-associated (yellow arrows).

**Figure 3.**
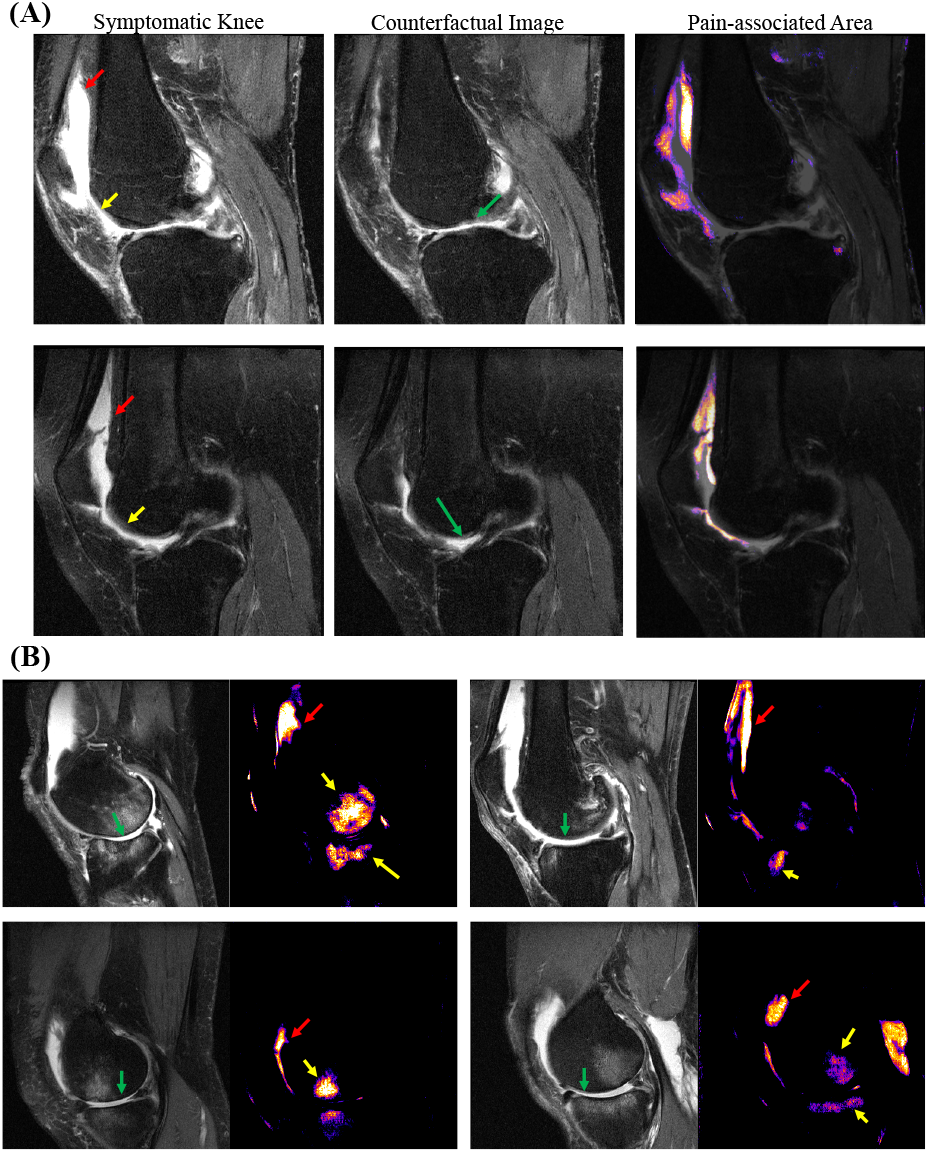
**The generative model was able to differentiate anatomical subregions in details and altered exclusively on the subregions associated with pain. (A) shows the comparison between the original image, counterfactuals, and the identified pain-associated area. The model identified effusion-synovitis in the patellar region (red arrows) and infrapatellar fat pad (yellow arrows) as pain-associated. Conversely, the effusion-synovitis in the central area and adjacent to the anterior cruciate ligament, not deemed pain-associated (green arrows), remains present in the images. (B) The model showed the capacity to characterize distinct source of pain without prior knowledge of anatomical structures. It identified effusion-synovitis in the patellar regions as pain-associated (red arrows) alongside bone marrow lesions (yellow arrows). Contrarily, the effusion-synovitis in the central region (green arrows) was not regarded as pain-associated, even when situated adjacent to bone marrow lesions and exhibiting lower intensity values.**

### 3.3 Distributions of source and location of the pain-associated regions identified

To explore the differential associations between subregions of effusion-synovitis and knee pain symptoms, we assessed the pain-associated effusion-synovitis volumes within each subregion and contrasted them with the overall volume. A significantly greater disparity in pain-associated effusion-synovitis volumes than in overall volume was identified by comparing symptomatic knees to their contralateral asymptomatic counterparts. Given that mild synovitis-effusion was prevalent both in asymptomatic and symptomatic knees, asymptomatic knees displayed considerable overall effusion-synovitis volumes in all the studied subregions, (28.0%, 60.0%, and 58.6% in patellar, central, and posterior subregions respectively, **Table 1**) in comparison to symptomatic knees. Conversely, knees without pain symptoms displayed markedly reduced pain-associated effusion-synovitis volumes relative to those with pain symptoms (8.9%, 15.3%, and 27.5% in patellar, central, and posterior subregions, respectively). These findings revealed that pain-associated volume of effusion-synovitis was a more distinctive biomarker for distinguishing knee with pain symptoms than the overall volume.

**Table 1.** Ratio of total and pain-associated effusion-synovitis volume in asymptomatic knees over symptomatic knees in the patellar, central and posterior subregion.

|  |  | **Patellar** | **Central** | **Posterior** |
| --- | --- | --- | --- | --- |
| **Volumetric ratio of asymptomatic / symptomatic knees** | Total Volume | 0.280 | 0.600 | 0.586 |
|  | Pain-associated Volume | 0.089 | 0.153 | 0.275 |

The most substantial pain-associated effusion-synovitis volume was found localized within the patellar subregion. In knees with pain symptoms, pain-associated effusion-synovitis in the patellar subregion constituted a substantial portion (39.2%, **Figure 4A**) and were highly correlated (0.934, **Figure 4B**) with the overall volume of effusion-synovitis. In contrast, markedly smaller percentages of overall effusion-synovitis in the central and posterior subregions were identified as pain-associated (14.2% and 15.6%, respectively), and the pain-associated volumes were merely moderately correlated to the overall volumes of effusion-synovitis in these subregions. This highlighted that pain-related effusion-synovitis as a more specific measurement for determining the contribution of effusion-synovitis subregions to pain symptoms, while the overall volumes were not adequately representative of the effusion-synovitis contribution to pain symptoms in the central and posterior subregions.

**Figure 4.**
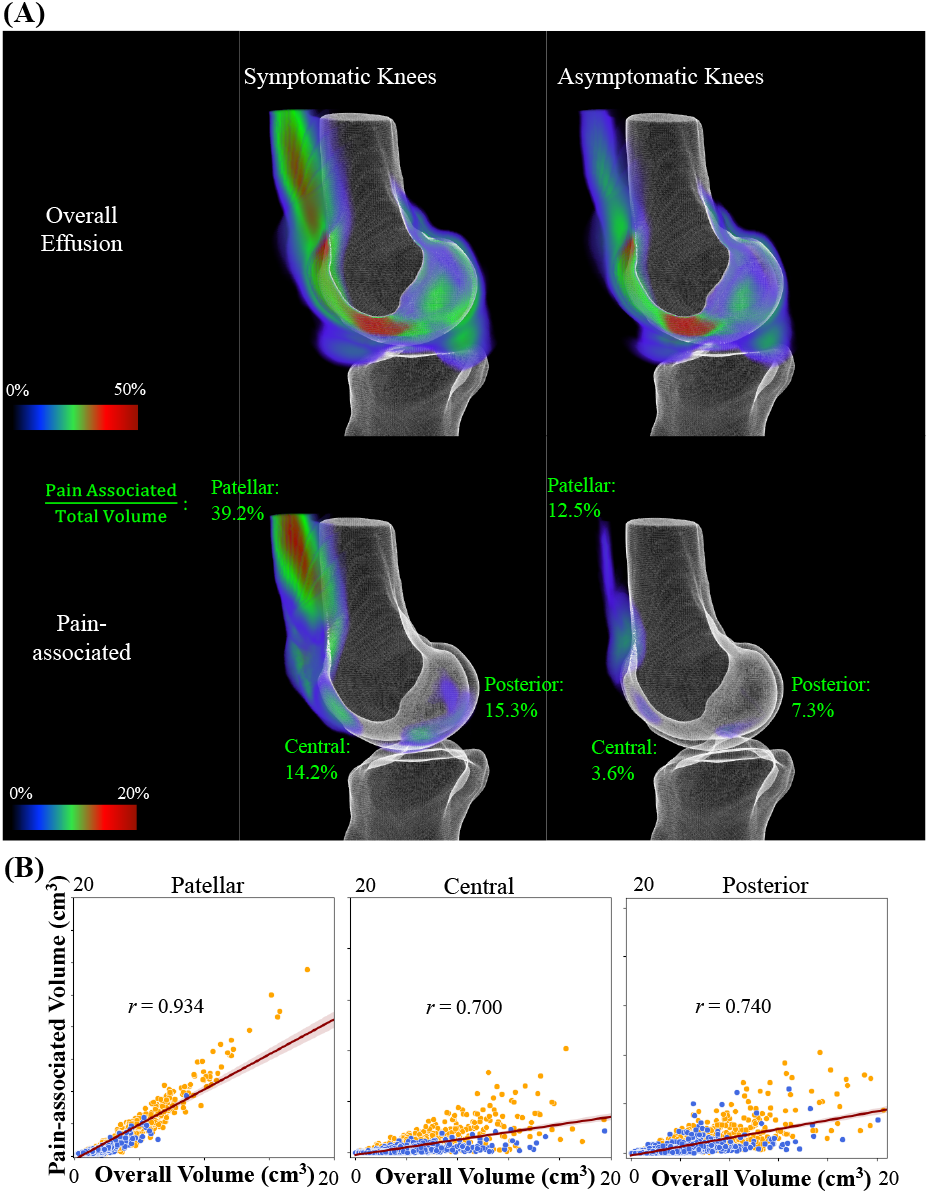
**(A) Spatial distribution of overall and pain-associated effusion-synovitis for symptomatic and asymptomatic knees. Green fonts indicate the portion of effusion-synovitis considered pain-associated in the patellar, central and posterior subregion. (B) Joint distribution of pain-associated and overall volume of effusion-synovitis for symptomatic and asymptomatic knees in different subregions.**

### 3.4 Clinical relevance of identified pain-associated areas

We evaluated the clinical relevance of the identified pain-associated areas by calculating the ORs of developing pain symptoms by severities of effusion-synovitis in each individual subregion, which were characterized based on their volumes and stratified into mild, moderate and severe categories. A significant correlation between pain symptoms and either moderate or severe pain-associated effusion-synovitis was observed, with the highest odds ratio found in knees exhibiting severe (4th quartile) pain-associated effusion-synovitis in the patellar region (OR 4.62, **Table 2**). Interestingly, the pain-associated volume of effusion-synovitis demonstrated a better association with pain symptoms in the subregions of central and posterior, as opposed to the overall volume. Knees with severe pain-associated effusion-synovitis in the central and posterior subregion exhibited higher odd ratios of experiencing pain symptoms compared to those with severe overall effusion-synovitis volume (OR 4.53 versus 2.34 in the central region, and OR 3.09 versus 2.74 in the posterior region). Furthermore, mild pain-associated effusion-synovitis was found to be associated with pain symptoms, whereas mild overall effusion-synovitis was not. This finding suggests that the pain-associated effusion-synovitis volume may serve as a more sensitive and reliable indicator of pain symptoms in specific subregions.

**Table 2.** Association of pain-associated effusion-synovitis with pain symptoms. OR = odds ratio; 95% CI = 95% confidence interval. *P<0.05.

| Abnormalities |  | Mild (Quartile 2) | Moderate (Quartile 3) | Severe (Quartile 4) |
| --- | --- | --- | --- | --- |
|  |  | OR (95% CI) | OR (95% CI) | OR (95% CI) |
| Effusion-synovitis, Patellar | Total Volume | 1.17 (0.99-1.39) | 1.45 (1.23-1.72)* | 4.87 (4.05-5.84)* |
|  | Pain-associated | 1.14 (0.96-1.36) | 1.23 (1.04-1.46)* | 4.62 (3.85-5.54)* |
| Effusion-synovitis, Central | Total Volume | 1.12 (0.94-1.32) | 1.44 (1.22-1.71)* | 2.34 (1.97-2.78)* |
|  | Pain-associated | 1.24 (1.05-1.48)* | 1.54 (1.30-1.82)* | 4.53 (3.78-5.43)* |
| Effusion-synovitis, Posterior | Total Volume | 1.17 (0.99-1.39) | 1.57 (1.32-1.86)* | 2.74 (2.31-3.26)* |
|  | Pain-associated | 1.26 (1.06-1.49)* | 1.53 (1.29-1.81)* | 3.09 (2.60-3.68)* |

Finally, to investigate the degree of informativeness of the identified pain-associated effusion-synovitis, we compared the ability to exclusively use pain-associated effusion-synovitis as the only input to predict which knee had unilateral knee pain and compared the results to those of the models using full MRI. We utilized the Siamese-neural network structure previously used for the prediction of unilateral knee pain symptoms (37) and the model performance of area under the ROC curve (AUROC) was calculated using five-fold cross-validation. It was found that using the information of pain-associated effusion-synovitis exclusively, the models could predict frequent knee pain in patients with unilateral knee pain with a performance comparable to that of the models using information from the entire MRI (**Figure 5**).

**Figure 5.**
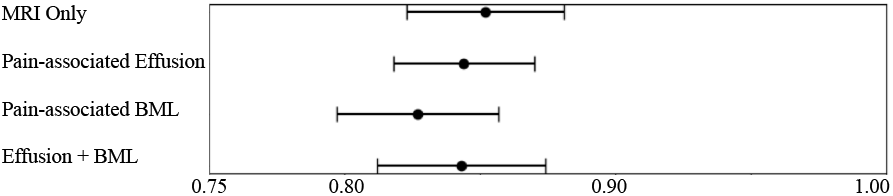
Model performance in predicting pain symptoms using magnetic resonance imaging (MRI), pain-associated effusion-synovitis, and BML exclusively.

## DISCUSSION

In this study, we utilized deep learning-based models to automatically characterize and quantify knee MRI inflammatory features associated with pain symptoms. To the best of our knowledge, this is the first such study. Traditionally, identifying imaging findings from knee MRI images has been a particularly difficult and time-consuming process, and reading times can be further increased in knees with extensive OA lesions (38). The challenge of associating imaging findings with pain symptoms is further compounded by the fact that MRI features in individual subregions may contribute to pain symptoms differently (34,39,40). Our algorithms allowed us to directly compare knee pairs with unilateral pain symptoms and enabled us to automatically characterize and quantify pain-associated effusion-synovitis, which have been shown to have a stronger correlation with pain symptoms than other MRI features such as cartilage defects (12). To investigate how different subregions of effusion-synovitis may contribute to OA symptoms, we quantified the volume of pain-associated effusion-synovitis, defined as the difference between the overall volume of effusion-synovitis in the original MR images and those in the generated counterfactuals. Our study found that knees with severe pain-associated effusion-synovitis in the patellar region had the highest odds ratio for significant pain symptoms compared to knees with significant effusion-synovitis in other studied subregions. This finding aligns with previous observations in which the subregions closest to the patella were most frequently involved in the pattern of synovitis, which was most correlated with pain (39).

The results of the present study indicate that pain-associated effusion-synovitis serves as a more sensitive and reliable indicators of pain symptoms than the overall volume of effusion-synovitis. In accordance with previous literature, only significant (moderate or advanced) effusion-synovitis volumes were found to be associated with significant knee pain symptoms (41). Small effusions, prevalent in asymptomatic knees and not considered clinically meaningful, posed challenges in their accurate definition and reliable exclusion from pain analysis (42). Our present approach therefore provided an objective way for distinguishing between effusion areas related to pain and those without such a correlation, as evidenced by the minimal presence of pain-associated effusion-synovitis in asymptomatic knees. Interestingly, small volume of pain-associated effusion-synovitis in the central and posterior areas were found to be associated with pain symptoms, while these associations were not found for knees with small volumes of overall effusion-synovitis. These results suggested that the relation between pain symptoms and the volumes of effusion-synovitis is nonlinear and location-dependent, wherein the presence and severity of pain-associated effusion-synovitis serve as more revealing predictors of pain than the overall effusion-synovitis volume.

The identification of pain-associated image features from magnetic resonance (MR) images necessitates high resolution of the detected regions to ensure accurate characterization among various anatomical subregions. In comparison to heatmap-based visualization techniques, such as Grad-CAM, our current approach offers markedly superior resolution, preserving the anatomical structures in the original images within the generated counterfactuals, thereby facilitating the differentiation of pain-associated lesions from different sources. As depicted in Figure 3B, the model was capable of identifying bone marrow lesions as pain-associated while excluding adjacent effusion-synovitis in the central subregion, thus demonstrating its ability to discern lesions of both large and small extents. More significantly, the model exhibited the capacity to distinguish different sources of pain without requiring explicit anatomical information as input. In essence, the model appeared to learn the criteria for characterizing lesions by source and extent, as well as determining the ways pain symptoms could be attributed to these lesions concurrently. This current model paves the way for future research on the relationship between lesions and pain symptoms without the need for predefined criteria for MRI scorings. Given that the detected pain-associated regions were defined at the pixel level, this model also facilitates future studies of lesion progression and its association with pain symptoms. This unbiased approach can aid in the discovery of novel imaging biomarkers correlated with pain symptoms.

While the present model demonstrates promising results for identifying pain-associated from MRI images of knees with OA, there is room for improvement. The cross-sectional nature of our study design limited our ability to examine time-varying factors related to pain symptoms. Our research incorporated instances from the OAI and follow-up cohorts, ensuring that knee pairs exhibiting significant pain differences at multiple time points were well-represented in the study. Nevertheless, we did not restrict our investigation to participants who consistently reported knee pain across various time points. It is worth noting that a considerable number of individuals with knee OA experience fluctuating pain symptoms (43). Although focusing exclusively on those with persistent pain might enhance the model’s accuracy in predicting pain symptoms, it could also result in a reduced training dataset, potentially undermining the generative model’s ability to detect abnormalities. In our study, we instead opted to exclude knee pairs with smaller magnitude differences in pain symptoms, as more pronounced pathology is expected to correlate with more significant differences in pain symptoms. Balancing these considerations is essential for optimizing the model’s overall performance in both predicting pain symptoms and identifying abnormalities.

In conclusion, our study demonstrated the ability of deep generative networks to automatically detect pain-associated abnormalities from MR images of knees with pain symptoms. Such algorithms have the potential to leverage large cohorts of image data and to allow the study of the associations between individual lesions and pain symptoms in knees with OA in different subregions, as demonstrated in our findings with regards to the relationship between effusion-synovitis and pain symptoms in individual subregions. Our findings can assist in identifying the source and location of OA-induced pain symptoms and can be beneficial in designing targeted and individualized treatment regimens.

## REFERENCE

1. Cross M, Smith E, Hoy D, Nolte S, Ackerman I, Fransen M, et al. The global burden of hip and knee osteoarthritis: estimates from the Global Burden of Disease 2010 study. Ann Rheum Dis 2014;73:1323–1330.

2. Neogi T. The epidemiology and impact of pain in osteoarthritis. Osteoarthritis Cartilage 2013;21:1145–1153.

3. Hunter DJ, Guermazi A, Roemer F, Zhang Y, Neogi T. Structural correlates of pain in joints with osteoarthritis. Osteoarthritis Cartilage 2013;21:1170–1178.

4. Bedson J, Croft PR. The discordance between clinical and radiographic knee osteoarthritis: A systematic search and summary of the literature. BMC Musculoskelet Disord 2008;9:116.

5. Finan PH, Buenaver LF, Bounds SC, Hussain S, Park RJ, Haque UJ, et al. Discordance between pain and radiographic severity in knee osteoarthritis: Findings from quantitative sensory testing of central sensitization. Arthritis Rheum 2013;65:363–372.

6. Dimitroulas T, Duarte R v., Behura A, Kitas GD, Raphael JH. Neuropathic pain in osteoarthritis: A review of pathophysiological mechanisms and implications for treatment. Semin Arthritis Rheum 2014;44:145–154.

7. Clauw DJ, Witter J. Pain and rheumatology: Thinking outside the joint. Arthritis Rheum 2009;60:321–324.

8. O’Neill TW, Felson DT. Mechanisms of Osteoarthritis (OA) Pain. Curr Osteoporos Rep 2018;16:611–616.

9. Neogi T, Felson D, Niu J, Nevitt M, Lewis CE, Aliabadi P, et al. Association between radiographic features of knee osteoarthritis and pain: Results from two cohort studies. BMJ (Online) 2009;339:498–501.

10. Chundru R, Baum T, Nardo L, Nevitt MC, Lynch J, McCulloch CE, et al. Focal knee lesions in knee pairs of asymptomatic and symptomatic subjects with OA risk factors - Data from the Osteoarthritis Initiative. Eur J Radiol 2013;82.

11. Guermazi A, Zaim S, Taouli B, Miaux Y, Peterfy CG, Genant HK. MR findings in knee osteoarthritis. Eur Radiol 2003;13:1370–1386.

12. Wenham CY, Conaghan PG. Imaging the painful osteoarthritic knee joint: what have we learned? Nat Rev Rheumatol 2009;5:149–158.

13. Hunter DJ, Guermazi A, Lo GH, Grainger AJ, Conaghan PG, Boudreau RM, et al. Evolution of semi-quantitative whole joint assessment of knee OA: MOAKS (MRI Osteoarthritis Knee Score). Osteoarthritis Cartilage 2011;19:990–1002.

14. Hunter DJ, Lo GH, Gale D, Grainger AJ, Guermazi A, Conaghan PG. The reliability of a new scoring system for knee osteoarthritis MRI and the validity of bone marrow lesion assessment: BLOKS (Boston–Leeds Osteoarthritis Knee Score). Ann Rheum Dis 2008;67:206–211.

15. Peterfy CG, Guermazi A, Zaim S, Tirman PFJ, Miaux Y, White D, et al. Whole-Organ Magnetic Resonance Imaging Score (WORMS) of the knee in osteoarthritis. Osteoarthritis Cartilage 2004;12:177–190.

16. Felson DT, Lynch J, Guermazi A, Roemer FW, Niu J, McAlindon T, et al. Comparison of BLOKS and WORMS scoring systems part II. Longitudinal assessment of knee MRIs for osteoarthritis and suggested approach based on their performance: data from the Osteoarthritis Initiative. Osteoarthritis Cartilage 2010;18:1402–1407.

17. Felson DT, Chaisson CE, Hill CL, Totterman SMS, Gale ME, Skinner KM, et al. The Association of Bone Marrow Lesions with Pain in Knee Osteoarthritis. Ann Intern Med 2001;134:541.

18. Felson DT, Niu J, Guermazi A, Roemer F, Aliabadi P, Clancy M, et al. Correlation of the development of knee pain with enlarging bone marrow lesions on magnetic resonance imaging. Arthritis Rheum 2007;56:2986–2992.

19. Hill CL, Hunter DJ, Niu J, Clancy M, Guermazi A, Genant H, et al. Synovitis detected on magnetic resonance imaging and its relation to pain and cartilage loss in knee osteoarthritis. Ann Rheum Dis 2007;66:1599–1603.

20. LeCun Y, Bengio Y, Hinton G. Deep learning. Nature 2015;521:436–444.

21. Litjens G, Kooi T, Bejnordi BE, Setio AAA, Ciompi F, Ghafoorian M, et al. A survey on deep learning in medical image analysis. Med Image Anal 2017;42:60–88.

22. Bayramoglu N, Nieminen MT, Saarakkala S. Automated detection of patellofemoral osteoarthritis from knee lateral view radiographs using deep learning: data from the Multicenter Osteoarthritis Study (MOST). Osteoarthritis Cartilage 2021;29:1432–1447.

23. Joseph GB, McCulloch CE, Nevitt MC, Link TM, Sohn JH. Machine learning to predict incident radiographic knee osteoarthritis over 8 Years using combined MR imaging features, demographics, and clinical factors: data from the Osteoarthritis Initiative. Osteoarthritis Cartilage 2022;30:270–279.

24. Guan B, Liu F, Haj-Mirzaian A, Demehri S, Samsonov A, Neogi T, et al. Deep learning risk assessment models for predicting progression of radiographic medial joint space loss over a 48-MONTH follow-up period. Osteoarthritis Cartilage 2020;28:428–437.

25. Lin T, Peng S, Lu S, Fu S, Zeng D, Li J, et al. Prediction of knee pain improvement over two years for knee osteoarthritis using a dynamic nomogram based on MRI-derived radiomics: a proof-of-concept study. Osteoarthritis Cartilage 2022.

26. Pedoia V, Lee J, Norman B, Link TM, Majumdar S. Diagnosing osteoarthritis from T2 maps using deep learning: an analysis of the entire Osteoarthritis Initiative baseline cohort. Osteoarthritis Cartilage 2019;27:1002–1010.

27. Hirvasniemi J, Runhaar J, Heijden RA van der, Zokaeinikoo M, Yang M, Li X, et al. The KNee OsteoArthritis Prediction (KNOAP2020) challenge: An image analysis challenge to predict incident symptomatic radiographic knee osteoarthritis from MRI and X-ray images. Osteoarthritis Cartilage 2022.

28. Tack A, Mukhopadhyay A, Zachow S. Knee menisci segmentation using convolutional neural networks: data from the Osteoarthritis Initiative. Osteoarthritis Cartilage 2018;26:680–688.

29. Xia X, Pan X, Li N, He X, Ma L, Zhang X, et al. GAN-based anomaly detection: A review. Neurocomputing 2022;493:497–535.

30. Yi X, Walia E, Babyn P. Generative adversarial network in medical imaging: A review. Med Image Anal 2019;58:101552.

31. Lemay A, Hoebel K, Bridge CP, Befano B, Sanjosé S de, Egemen D, et al. Improving the repeatability of deep learning models with Monte Carlo dropout. NPJ Digit Med 2022;5:174.

32. Nguyen D, Sadeghnejad Barkousaraie A, Bohara G, Balagopal A, McBeth R, Lin M-H, et al. A comparison of Monte Carlo dropout and bootstrap aggregation on the performance and uncertainty estimation in radiation therapy dose prediction with deep learning neural networks. Phys Med Biol 2021;66:054002.

33. Ronneberger O, Fischer P, Brox T. U-Net: Convolutional Networks for Biomedical Image Segmentation. 2015.

34. Wang X, Blizzard L, Halliday A, Han W, Jin X, Cicuttini F, et al. Association between MRI-detected knee joint regional effusion-synovitis and structural changes in older adults: a cohort study. Ann Rheum Dis 2016;75:519–525.

35. Guan H, Liu M. Domain Adaptation for Medical Image Analysis: A Survey. IEEE Trans Biomed Eng 2022;69:1173–1185.

36. Chang GH, Park LK, L. NA, Jhun RS, Surendran T, Lai J, et al. Subchondral Bone Length in Knee Osteoarthritis: A Deep Learning–Derived Imaging Measure and Its Association With Radiographic and Clinical Outcomes. Arthritis & Rheumatology 2021;73:2240–2248.

37. Chang GH, Felson DT, Qiu S, Guermazi A, Capellini TD, Kolachalama VB. Assessment of knee pain from MR imaging using a convolutional Siamese network. Eur Radiol 2020;30:3538–3548.

38. Jaremko JL, Jeffery D, Buller M, Wichuk S, Mcdougall D, Lambert RG, et al. Preliminary validation of the Knee Inflammation MRI Scoring System (KIMRISS) for grading bone marrow lesions in osteoarthritis of the knee: data from the Osteoarthritis Initiative. Open 2017;3:355. Available at: https://www.clinicaltrials.gov/ct2/show/NCT00686439.

39. Lange-Brokaar BJE de, Ioan-Facsinay A, Yusuf E, Visser AW, Kroon HM, Osch GJVM van, et al. Association of Pain in Knee Osteoarthritis With Distinct Patterns of Synovitis. Arthritis & Rheumatology 2015;67:733–740.

40. Aso K, Shahtaheri SM, McWilliams DF, Walsh DA. Association of subchondral bone marrow lesion localization with weight-bearing pain in people with knee osteoarthritis: data from the Osteoarthritis Initiative. Arthritis Res Ther 2021;23:35.

41. Guermazi A, Roemer FW, Hayashi D, Crema MD, Niu J, Zhang Y, et al. Assessment of synovitis with contrast-enhanced MRI using a whole-joint semiquantitative scoring system in people with, or at high risk of, knee osteoarthritis: the MOST study. Ann Rheum Dis 2011;70:805–811.

42. Boks SS, Vroegindeweij D, Koes BW, Hunink MMGM, Bierma-Zeinstra SMA. Magnetic Resonance Imaging Abnormalities in Symptomatic and Contralateral Knees. Am J Sports Med 2006;34:1984–1991.

43. Hawker GA, Stewart L, French MR, Cibere J, Jordan JM, March L, et al. Understanding the pain experience in hip and knee osteoarthritis – an OARSI/OMERACT initiative. Osteoarthritis Cartilage 2008;16:415–422.

